# Primary cilium-dependent cAMP/PKA signaling at the centrosome regulates neuronal migration

**DOI:** 10.1101/765925

**Authors:** Julie Stoufflet, Maxime Chaulet, Mohamed Doulazmi, Coralie Fouquet, Caroline Dubacq, Christine Métin, Alain Trembleau, Pierre Vincent, Isabelle Caillé

## Abstract

The primary cilium (PC) is a small centrosome-assembled organelle, protruding from the surface of most eukaryotic cells. It plays a key role in cell migration, but the underlying mechanisms are unknown. Here, we show that the PC regulates neuronal migration via cAMP production activating centrosomal Protein Kinase A (PKA). Biosensor live-imaging revealed a periodic cAMP hotspot at the centrosome of embryonic, postnatal and adult migrating neurons. Genetic ablation of the PC, or knock-down of ciliary Adenylate Cyclase 3, caused hotspot disappearance and migratory defects, with defective centrosome/nucleus coupling and altered nucleokinesis. Delocalization of PKA from the centrosome phenocopied the migratory defects. Our results show that the PC and centrosome form a single cAMP-signaling unit dynamically regulating migration, further highlighting the centrosome as a signaling hub.

The primary cilium regulates neuronal migration via cyclic AMP production activating Protein Kinase A at the centrosome

---

The PC is a small microtubule-based organelle templated by the centrosome and involved in multiple cellular events, including cell motility (*1*) and neuronal migration in particular (*2*–*5*). How the PC regulates neuronal migration is largely unknown.

Neuronal migration is essential to the formation of functional neural circuits, with defective migration leading to severe brain malformations and involved in psychiatric disorders (*6*). Migrating neurons display a cyclic saltatory migration with alternations of somal translocation and pauses. Nucleus and centrosome move forward in a “two-stroke” cycle, with the centrosome moving first within a swelling in the leading process (centrokinesis, CK) and the nucleus following subsequently (nucleokinesis, NK) (Fig. 1A). Centrosome dynamics is thus pivotal in the regulation of migration, through microtubular centrosome to nucleus coupling allowing NK (*7*–*9*).

**Figure 1:**
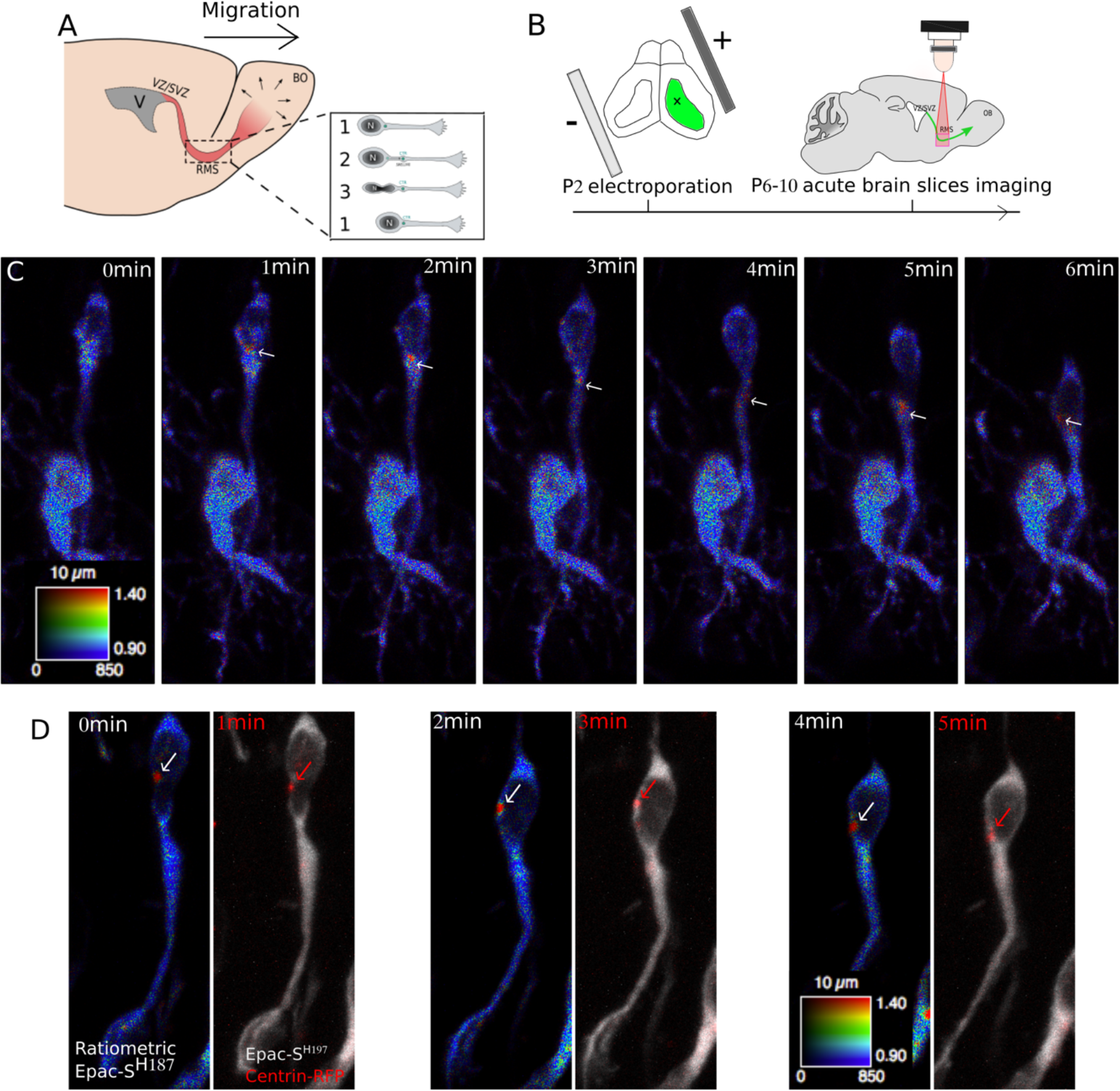
cAMP dynamics in migrating neurons of the postnatal RMS. A) SVZ-OB migration pathway and cyclic saltatory migration. 1: pause. 2: leading process extension and centrosome forward movement CK. 3: NK. B) Postnatal experimental procedure. C) Live two-photon imaging of a representative migrating neuron transfected by Epac-S^H187^. The white arrows show a dynamic cAMP hotspot present during NK. A non-migrating transfected cell is present in the field of view. Scale-bar: 10μm D) Live two-photon and confocal imaging of a representative migrating neuron transfected by Epac-S^H187^ and Centrin-RFP. A dynamic cAMP hotspot (white arrow) is present at the centrosome (red arrow).

As the PC is a cAMP-rich source (*10*–*13*), we asked whether it could regulate neuronal migration through cAMP dynamics, in the context of migration along the postnatal Rostral Migratory Stream (RMS) from the ventricular/subventricular zone to olfactory bulb (OB) (Fig. 1A).

To investigate cAMP dynamics in migrating neurons, we electroporated neonate mice with an intraventricularly-injected plasmid to express the FRET (Förster Resonance Energy Transfer)-based cAMP-specific biosensor Epac-S^H187^ (Fig. 1B) (*14*). Biosensor-expressing neurons were live-imaged in acute sections of the RMS with a two-photon microscope. Ratiometric analysis revealed that a single cAMP-rich region (hereafter called hotspot) was present during NK and disappeared during pauses (Fig. 1C and movie 1). 82% of migrating neurons displayed a dynamic hotspot (101 neurons, 134 NK from 10 mice). This hotspot specifically reflected a cAMP local enrichment since it was absent from neurons transfected with a version of Epac-S^H187^ with mutated cAMP-binding site (movie 2). The cAMP-hotspot diameter was 965±95nm and the maximum hotspot ratio level was constant during NK, being 52% (±2%) higher than the mean cell ratio. The hotspot stereotypically appeared at the front of the cell body before moving into the leading process, where it remained until after the nucleus performed NK. As this movement resembled the centrosome movement during migration, we co-electroporated the biosensor and centrin-RFP to label the centrosome. The hotspot co-localized with the centrosome before and during NK (Fig. 1D). We thus discovered a dynamic cAMP hotspot at the centrosome of RMS migrating neurons.

The PC is a small rod-shaped organelle linked to the centrosome by its axoneme (Fig. 2A). Embryonic migrating neurons assemble a PC (*2*,*3*,*15*) and immunostainings revealed that it is also the case for RMS migrating neurons (Fig. 2B, Fig. S1), similar to what was described recently (*5*). The PC is a cAMP-rich region (*10*–*13*), raising the possibility that the cAMP hotspot at the centrosome could be PC-dependent. To test this hypothesis, we used two mouse lines in which ablation of the PC can be genetically-induced by Cre recombination (Kif3a^lox/lox^ and Rpgrip1l^lox/lox^ mice). Cre-electroporated mice were compared to GFP-electroporated mice. Cre electroporation led to an efficient ablation of the PC in both lines (Fig. S1). Co-transfection of the biosensor with Cre in floxed mice showed that cilium ablation led to a concomitant disappearance of the hotspot in both lines (Fig. 2C and movies 3,4), showing that the PC is necessary for the cAMP hotspot formation.

**Figure 2:**
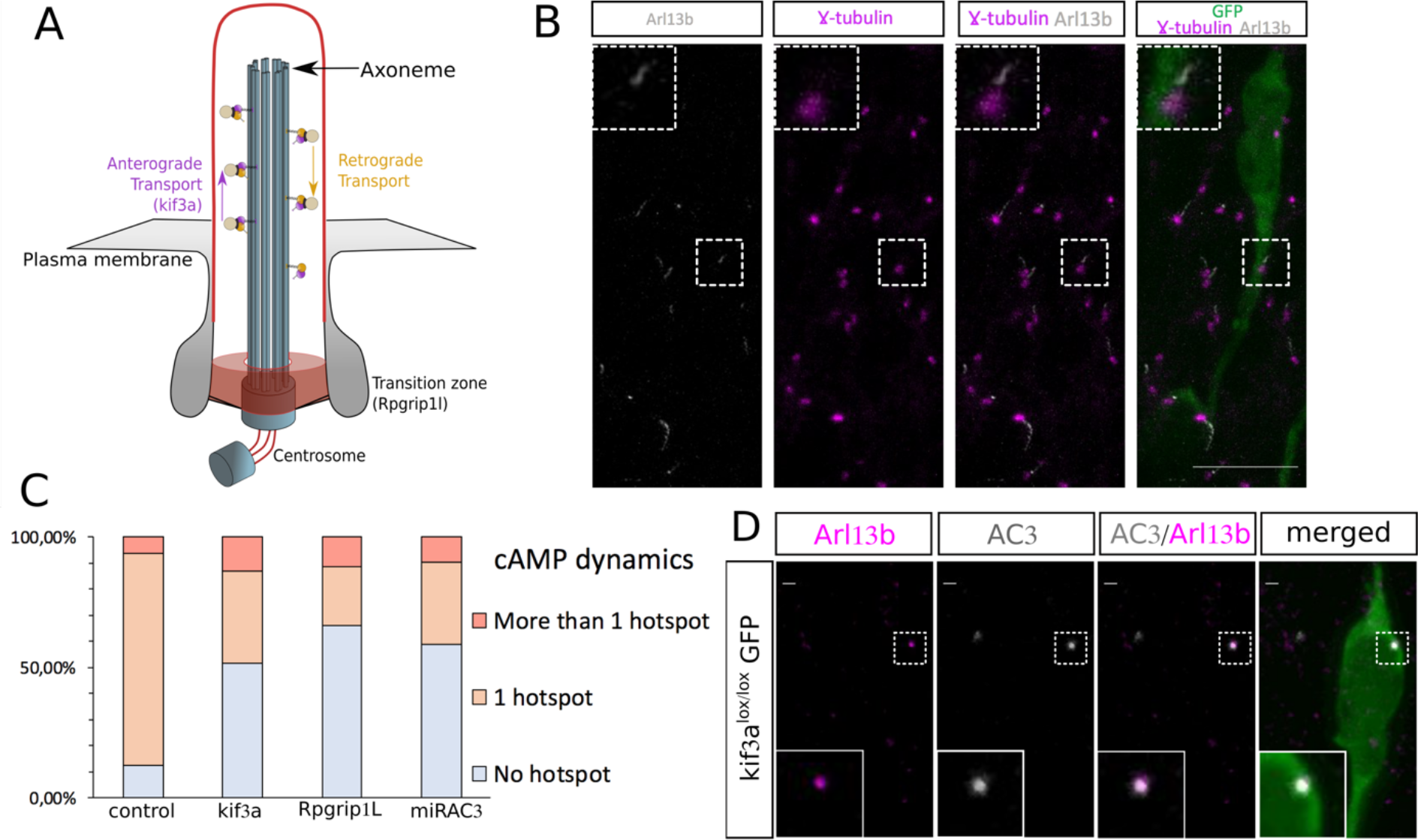
Primary cilium ablation and AC3 knock-down lead to disappearance of the cAMP-hotspot in the majority of neurons. A) Primary cilium. B) Immunohistochemistry of a GFP-positive RMS neuron showing a short Arl13b-positive PC (gray) connected to the γ-tubulin-positive centrosome (magenta). The PC length was in average 0,827±0.058μm (N=3 mice, n=50 PC). Scale-bar: 10μm C) Percentage of neurons with no hotspot, one or more than one hotspot in control or Kif3a^lox/lox^ and Rpgrip1^lox/lox^ CRE-transfected neurons. (Control 82% cells with hotspot (N=10 n=101) versus CRE-Kif3a^lox/lox^ 30% (N= 4 n=36) and CRE-Rpgrip1l^lox/lox^ 20% (N=7 n=52) and miRAC3 31% (N=3, n=62). (Fisher’s Exact Test, p<0.001) D) Immunocytochemistry of a GFP-positive neuron showing AC3 subcellular localization (magenta) in the Arl13B-positive PC (gray). Scale-bar: 1μm

The membrane-bound Adenylate Cyclase 3 (AC3) is the predominant cAMP-producer in neuronal PC (*16*). Immunostaining indeed revealed that AC3 is subcellularly localized in the PC (Fig. 2D). Co-electroporation of a microRNA targeting AC3 mRNA coupled to GFP (Fig. S2) with the biosensor showed that AC3 KD led to similar disappearance of the hotspot as PC ablation (movie 5, Fig. 2C), showing that ciliary AC3 produces the hotspot.

To assess whether the hotspot influences migration, we compared the migration of control neurons with the migration of cilium-ablated and AC3 knocked-down neurons, conditions leading to hotspot disappearance. Kif3a^lox/lox^ mice were thus transfected with GFP, Cre-GFP or miRAC3-GFP. Tracking of the GFP cells showed that cilium ablation and AC3 KD both slowed-down migration (Fig. 3A, movies 6-8) and increased pausing time (Fig. 3B). The frequency of NK was reduced (Fig. 3C), while the speed and distance of NK were unchanged. To confirm that the phenotype of recombined Kif3a^lox/lox^ mice was due to cilium ablation, we performed the same experiment in Rpgrip1l^lox/lox^ mice and observed the same defects (Fig. S3).

**Figure 3:**
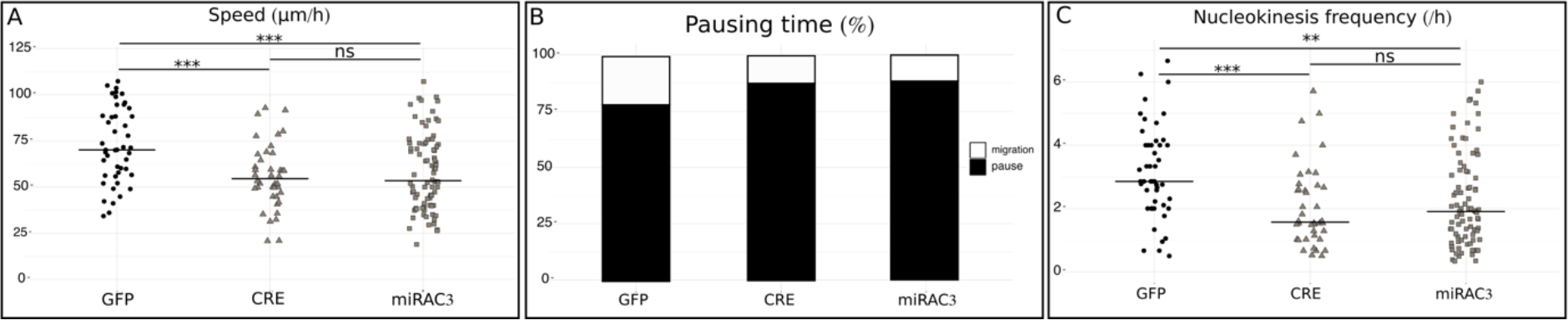
Migration defects after primary cilium ablation and AC3 knock-down. A) Speed of neurons electroporated with GFP, CRE-GFP or miRAC3-GFP in Kif3a^lox/lox^ background. GFP: 75.49±3.48μm/h; CRE: 55.81±2.74μm/h; miRAC3: 58.92±2.35 μm/h (one-way anova (F_(4,387)_=7.87, p<0.001, followed by Tukey-HSD (***p<0.001) Percentage of pausing time of neurons electroporated with GFP, CRE-GFP or miRAC3-GFP in Kif3a^lox/lox^ background. GFP: 76%; CRE: 87%; miRAC3: 84% (Pearson’s Chi-squared X^2^=40.032, p<0.001) C) NK frequency of neurons electroporated with GFP, CRE-GFP or miRAC3-GFP in Kif3a^lox/lox^ background. GFP: 3.15±0.21 NK/h; CRE: 2.06±0.20NK/h; miRAC3: 2.27±0.16NK/h (One-way Kruskal-Wallis (X2=19.544, p<0.001, df=4, followed by Nemenyi (**p<0.01, ***p<0.001) The black line represents the median. GFP: N=3, n=48, CRE: N=3, n=40, miRAC3: N=3, n=85

Given the centrosomal localization of the hotspot, we wondered whether the hotspot could directly play a role in centrosome dynamics. We co-transfected centrin-RFP either with GFP, Cre-GFP or miRAC3-GFP in Kif3a^lox/lox^ background and co-tracked the cell body and centrosome in migrating neurons (movie 9 (GFP) and 10 (CRE)). The maximum nucleus-centrosome distance reached by the centrosome during CK was reduced in cilium-ablated and AC3 knocked-down neurons, compared to GFP neurons (Fig. S4A). Furthermore, inefficient CK (CK not followed by NK) was increased in cilium-ablated and AC3 KD neurons compared to GFP neurons (Fig. S2B), suggesting that the hotspot is necessary for proper microtubular coupling of centrosome and nucleus.

Altogether, our results show that the ciliary-dependent cAMP-hotspot regulates migration by a direct action on the centrosome, with consequences on centrosome/nucleus coupling and NK.

PKA is localized at the centrosome of diverse cell types including neurons (*17*–*19*). Immunostaining for the catalytic subunit of PKA (cPKA) showed that it is the case in RMS migrating cells, where it appears as a diffuse area surrounding the centrosome (Fig. 4A), while it was not detected in the PC (not shown). To test the role of PKA, we electroporated the regulatory subunit of PKA devoid of its centrosomal anchoring domain (dominant-negative, dnPKA) coupled to GFP, which traps the endogenous cPKA in the cytoplasm (*19*) (Fig. 4B, Fig. S5). Similar to cilium ablation and AC3 KD, dnPKA slowed-down migration (Fig. 4C,), increased pausing phases (Fig. 4D) and reduced frequency of NK (Fig. 4E) (movies 6,11).

**Figure 4:**
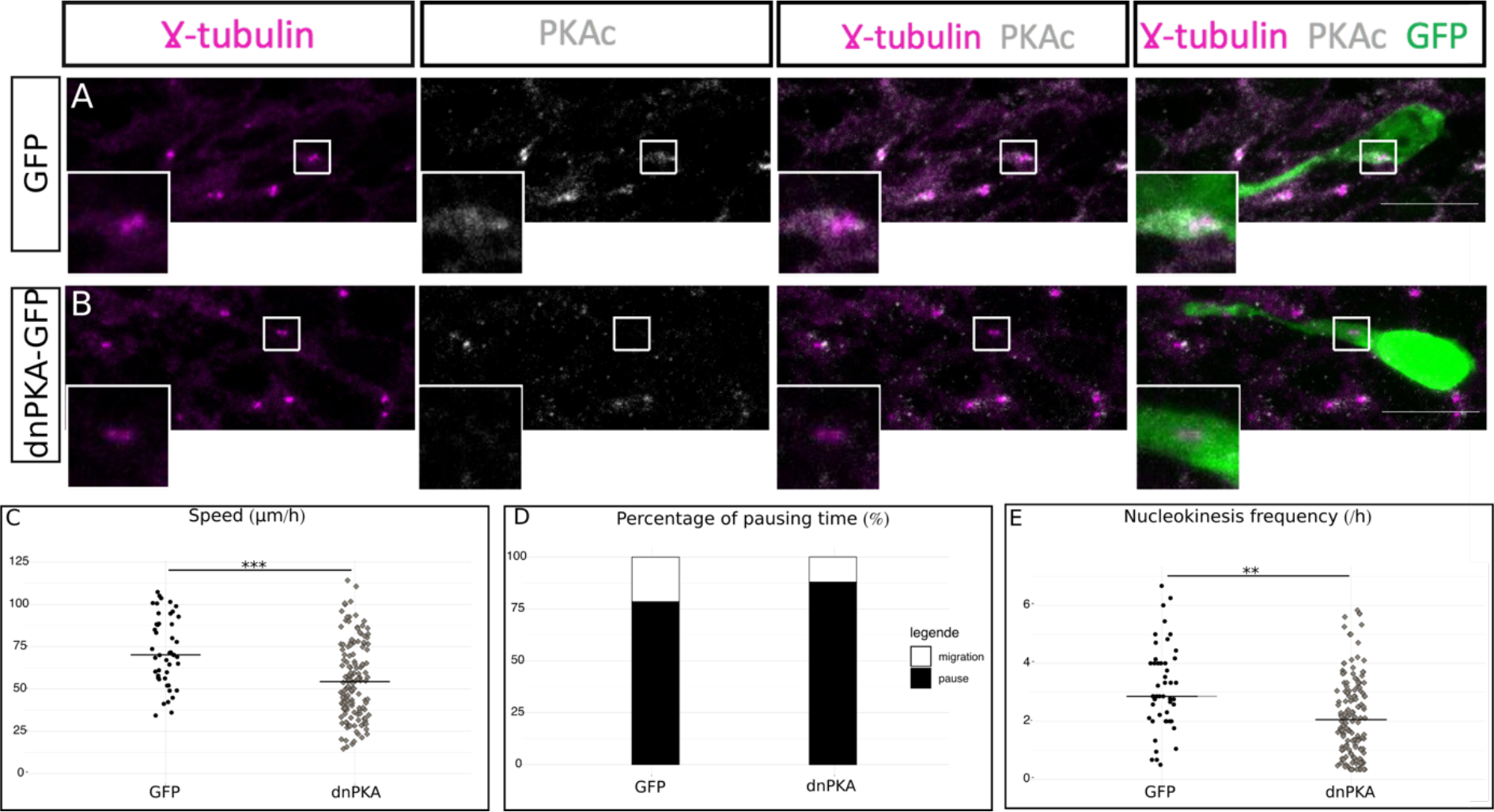
PKA is centrosomal and its delocalization phenocopies cilium ablation. A) Immunocytochemistry of a GFP-positive neuron showing PKAc subcellular localization (grey) around γ-tubulin-positive centrosome (magenta). Scale-bar: 10μm B) Immunocytochemistry of a dnPKA-GFP-positive neuron showing a delocalization of PKAc (grey) from the γ-tubulin-labeled centrosome (magenta). Scale-bar: 10μm. C) Speed of neurons electroporated with GFP or dnPKA-GFP in Kif3a^lox/lox^ background. GFP: 75.49±3.48μm/h; dnPKA 55.76±1.90μm/h (one-way anova (F_(4,383)_=7.87, p<0.001, followed by Tukey-HSD, ***p<0.001) D) Percentage of pausing time of neurons electroporated with GFP or dnPKA-GFP in Kif3a^lox/lox^ background. GFP: 76%; dnPKA: 86% (Pearson’s Chi-squared X^2^=67.25, p<0.001) E) NK frequency of neurons electroporated with GFP or dnPKA-GFP in Kif3a^lox/lox^ background. GFP 3.15±0.21 NK/h; dnPKA 2.23±0.12NK/h (One-way Kruskal-Wallis (X^2^=19.57, p<0.001, df=4, followed by Nemenyi (***p<0.001) The black line represents the median. GFP: N=3, n=48, dnPKA: N=3, n=146

Moreover, co-transfection with centrin-RFP revealed that centrosome/nucleus coupling was altered: the maximum distance between centrosome and nucleus during CK was reduced as well as inefficient CK (Fig. S6). Delocalization of PKA from the centrosome thus phenocopies cilium ablation and AC3 KD, suggesting that centrosomal PKA is the downstream effector of the hotspot. To confirm this result, we co-transfected Kif3a^lox/lox^ mice with Cre and dnPKA (Fig. S7): the non-additivity of the phenotypes shows that PKA indeed acts downstream of the ciliary-produced cAMP to regulate migration at the centrosome.

To test whether this pattern of cAMP dynamics is a general feature of migrating neurons, we assessed its presence in other types of neuronal migration. Radially-migrating neurons in the postnatal OB display a transient centrosomal hotspot (Fig. S8A, movie 13) as well as tangentially-migrating neurons of the adult RMS (Fig. S8B, movie 14) and radially-migrating neurons of the embryonic cortex (Fig. S8C, movie 15).

We can thus conclude that the centrosomal cAMP-hotspot is a general feature of migrating neurons, independent of age and type of migration.

The importance of the PC for neuronal migration has been inferred from ciliopathies
(*20*) and evidenced in embryonic cortical interneurons (*2*, *3*). The PC is often considered as a signal integrator, with paramount importance for Shh signaling (*2*, *21*). However, the mechanisms by which cilium-mediated signaling is converted into a migratory response are not understood (*22*). Our data provide such an intracellular mechanism, whereby the centrosome acts as a cAMP/PKA signaling platform downstream of the PC, with AC3-produced cAMP reaching the centrosome, where it locally activates PKA to regulate centrosome dynamics and NK. Very recently, the PC in postnatal and adult RMS was shown to cyclically emerge from the plasma membrane of migrating neurons before and during NK (*5*). Strikingly, this is coincident with the hotspot formation, so that the cyclic emergence of the PC could directly explain the cAMP hotspot cyclicity.

The centrosome as a signaling center is an attractive concept (*23*). Its importance as a cAMP signaling platform was hypothesized with the discovery of PKA centrosomal localization (*17*, *18*). Interestingly, a study using a centrosome-targeted FRET-biosensor in cultured cells reported a low cAMP microdomain at the centrosome, important for cell cycle progression (*24*). Regulation of the pattern of division of neural progenitors also involves PKA at the centrosome (*19*). Here, we report a role of the centrosome as a cAMP signaling platform in a non-mitotic cell, with a functional output on the regulation of migration through local PKA activation. The co-localization of the cAMP-hotspot with PKA at the centrosome might ensure that the concentration of cAMP locally exceeds the activation threshold of PKA (*25*).

To conclude, we show that the PC and centrosome are functionally linked by cAMP signaling in migrating neurons. We propose that PC and centrosome may be considered as a single cAMP-signaling unit, where the PC is a cAMP producer through local AC3, and the centrosome is the effector, through activation of centrosomal PKA by short-range diffusion of ciliary cAMP. This might create a direct link between the PC and the microtubule-organizer function of the centrosome. This cAMP dialogue between PC and centrosome might exist in other ciliated cell types to subserve diverse functions in health and disease.

## Supporting information

Movie 1: cAMP biosensor in control migrating neuron

Movie 2: cAMP insensitive biosensor in control migrating neuron

Movie 3: cAMP biosensor in Kif3a recombined migrating neuron

Movie 4: cAMP biosensor in RPGrip1L recombined migrating neuron

Movie 5: cAMP biosensor in AC3 knocked-down migrating neuron

Movie 6: GFP migrating neurons (control condition)

Movie 7: Cre-GFP migrating neurons (Kif3a recombined condition)

Movie 8: MirAC3-GFP migrating neurons (AC3 KD condition)

Movie 9: GFP + CentrinRFP migrating neuron (control condition)

Movie 10: Cre-GFP + CentrinRFP migrating neuron (Kif3a recombined condition)

Movie 11: dnPKA-GFP + tdTomato migrating neurons (delocalized PKA condition)

Movie 12: dnPKA-GFP + Cre-tdTomato migrating neurons (delocalized PKA + Kif3a recombined condition)

Movie 13: cAMP biosensor in radially migrating neuron of the postnatal olfactory bulb

Movie 14: cAMP biosensor in tangentially migrating neuron of the adult RMS

Movie 15: cAMP biosensor in radially migrating neuron of the embryonic cortex (E19)

---

**Figure S1.**
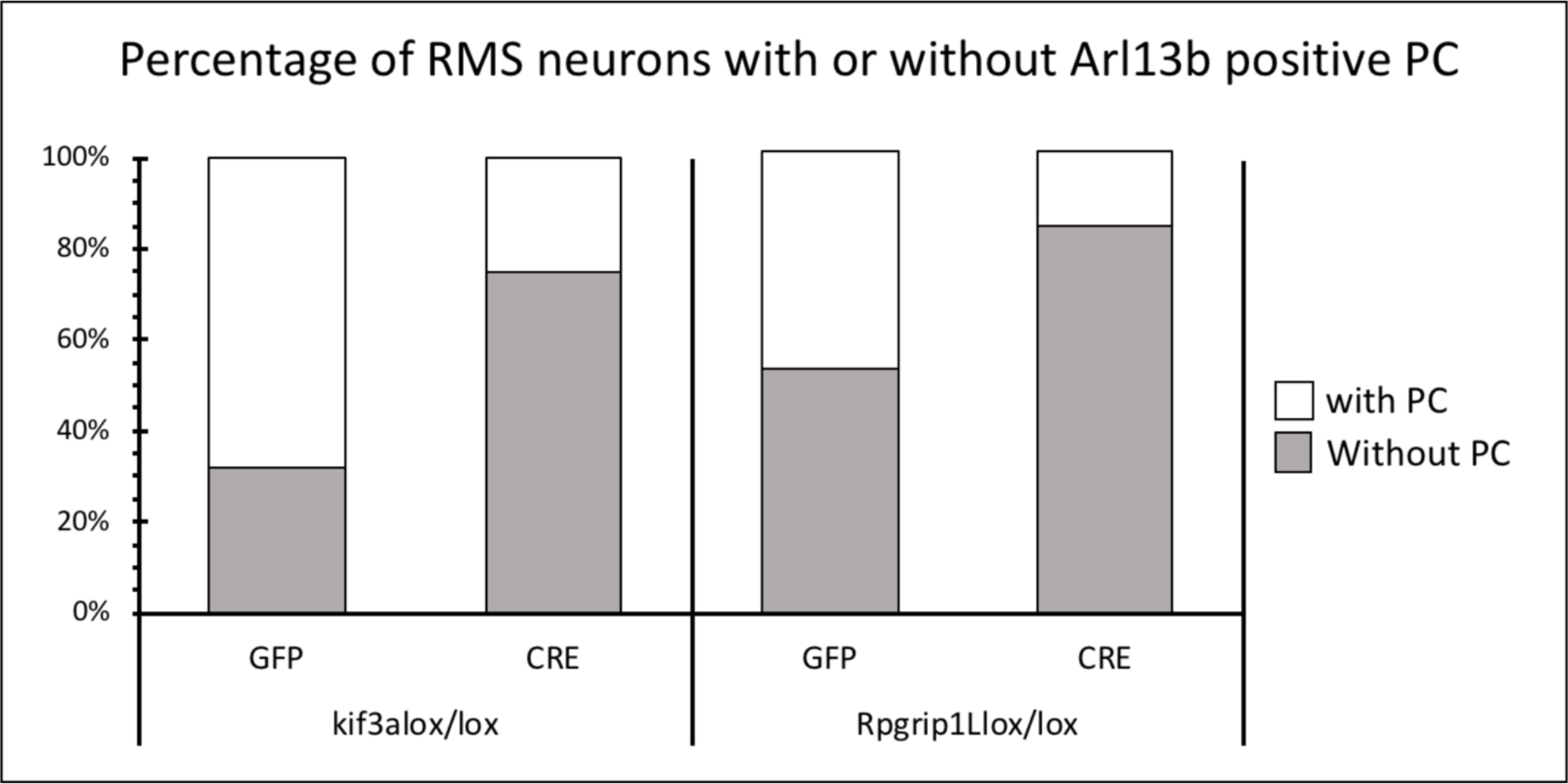
Efficiency of cilium ablation after Cre electroporation in Kif3a^lox/lox^ and Rpgrip1l^lox/lox^ mice: The percentage of GFP-positive neurons displaying a γ-tubulin positive centrosome associated with an Arl13B positive PC was significantly reduced in Cre-injected Kif3a^lox/lox^ mice (*26*) (25% in CRE (n=14, N=3) versus 68,3% in GFP (n=109, N=3), p<0.001) and Rpgrip1l^lox/lox^ mice (*27*) (16.2% in CRE (n=99, N=3) versus 47% in GFP (n=68, N=3), Chi-squared test p<0.001).

**Figure S2.**
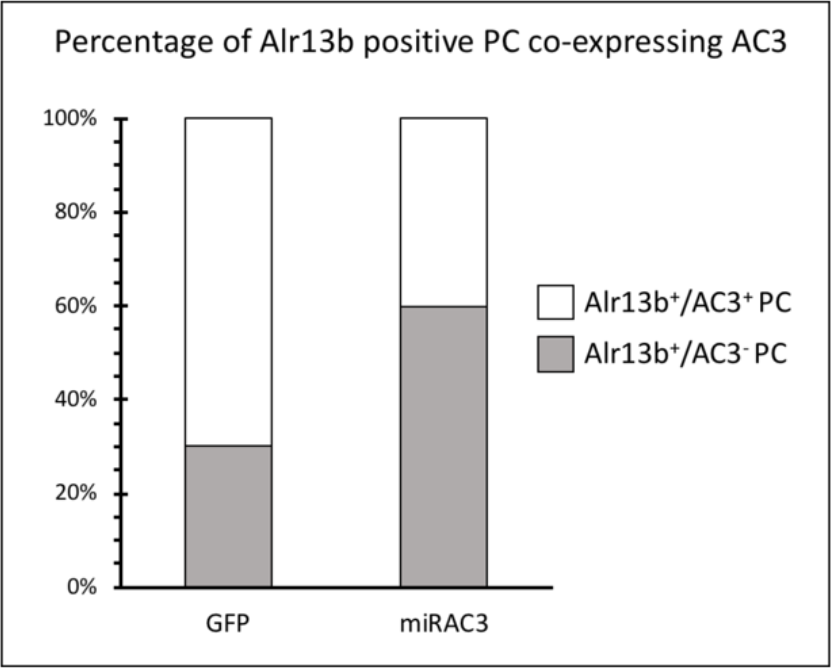
Efficiency of MirAC3 knock-down. The percentage of GFP neurons displaying an Arl13B and AC3 immunoreactive PC was significantly reduced in miRAC3 transfected neurons (70% in GFP (N=3, n=47) versus 40% in mirAC3 (N=3, n=63)). (X-squared=10.066, df=1, p<0.01)

**Figure S3:**
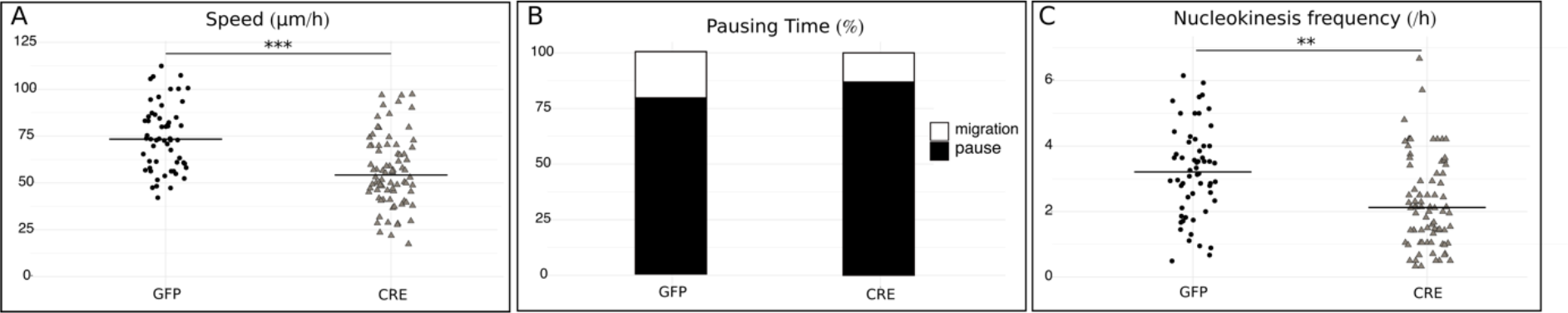
Migration defects after primary cilium ablation in Rpgrip1l^lox/lox^ mice. A) Speed of neurons electroporated with GFP or CRE-GFP in Rpgrip1l^lox/lox^ background: the speed was slowed-down in cilium-ablated neurons (GFP 75.33±2.45 μm/h versus CRE 57.88±2.43μm/h, p<0.001, student-test t(136) = 4.86, p<0.001) B) Percentage of pausing time of neurons electroporated with GFP or CRE-GFP in Rpgrip1L background: the pausing time was increased in cilium-ablated neurons (GFP 77% versus CRE 85%, p<0.001, Pearson’s Chi-squared test X^2^= 48.66, p-value < 0.001) C) Nuclear translocation frequency of neurons electroporated with GFP or CRE-GFP in Rpgrip1l background: the frequency of NK was reduced in cilium-ablated neurons (GFP 3.23± 0.18 NK/h versus CRE 2.26± 0.15 NK/h, p=0.03, Mann-Whitney test U (136) =3.01, p= 0.03) The black lines represent the median. GFP: N=4, n=58, CRE: N=3, n=80

**Figure S4:**
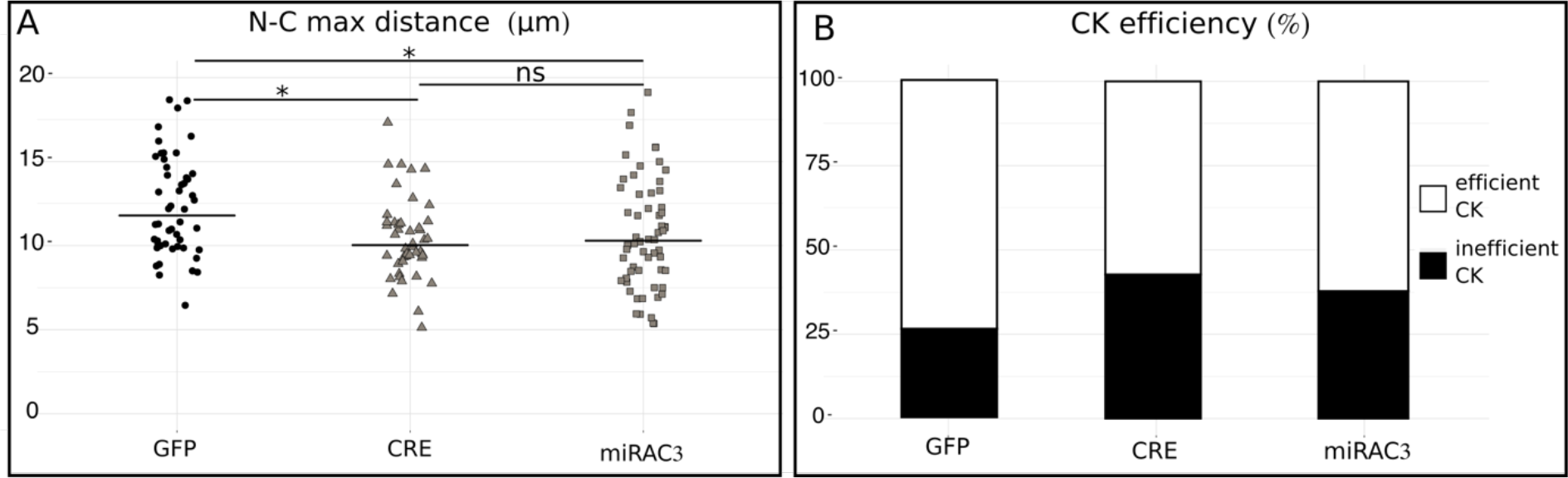
centrokinesis defects after cilium ablation or AC3KD. A) Maximal distance between nucleus and centrosome during CK of neurons electroporated with centrin-RFP and GFP, CRE-GFP or miRAC3-GFP in Kif3a^lox/lox^ background: the maximal distance was reduced in cilium-ablated and AC3 KD neurons (GFP 12.32±0.42μm versus CRE 10.42±0.35μm and miRAC310.84±0.44μm, One-way Kruskal-Wallis test (Chi square X= 13.54, p = 0.003, df = 3, followed by Nemenyi test (* p<0.05)) B) Percentage of CK efficiency in neurons electroporated with centrin-RFP and GFP, CRE-GFP or miRAC3-GFP in Kif3a^lox/lox^ background: the percentage of inefficient CK was increased in cilium-ablated and AC3 KD neurons. (GFP 26% versus Cre 43% and miRAC3 38%, Pearson’s Chi-squared test X= 10.75, p=0.01) The black lines represent the median. GFP: N=3, n=50, CRE: N=3, n=46, miRAC3: N=3, n=61

**Figure S5:**
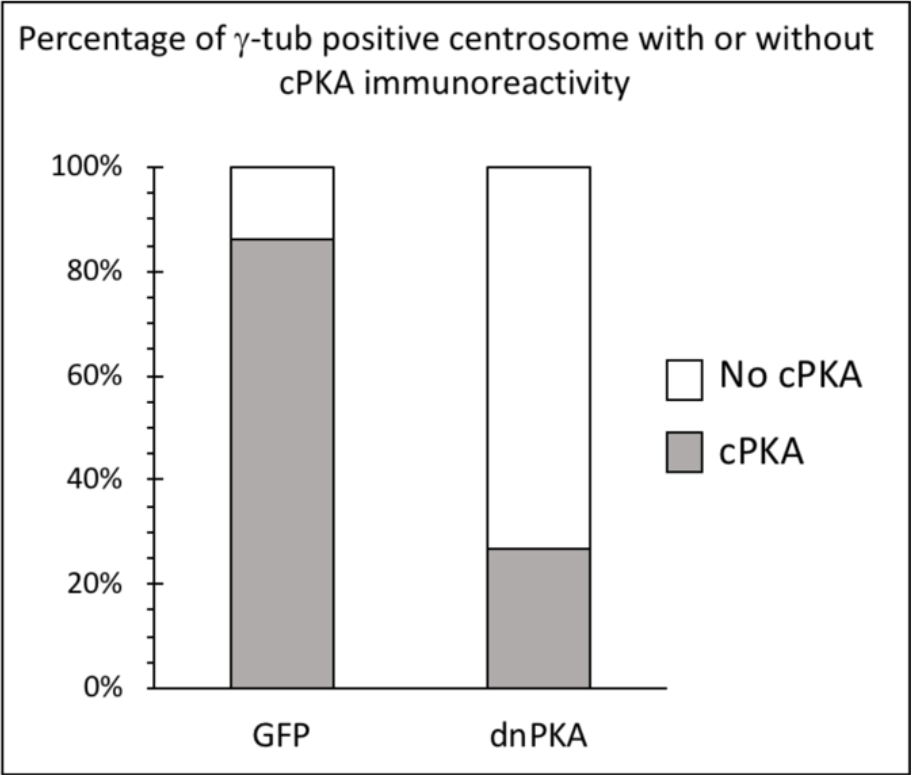
efficiency of PKA delocalization by dnPKA. dnPKA efficiently delocalized PKA from the centrosome (26.67% of neurons with visible cPKA immunoreactivity at the centrosome in dnPKA (n=30 N=2) versus 86.05% in GFP (n=44 N=3) (X-squared = 26.352, df=1, p<0.001).

**Figure S6:**
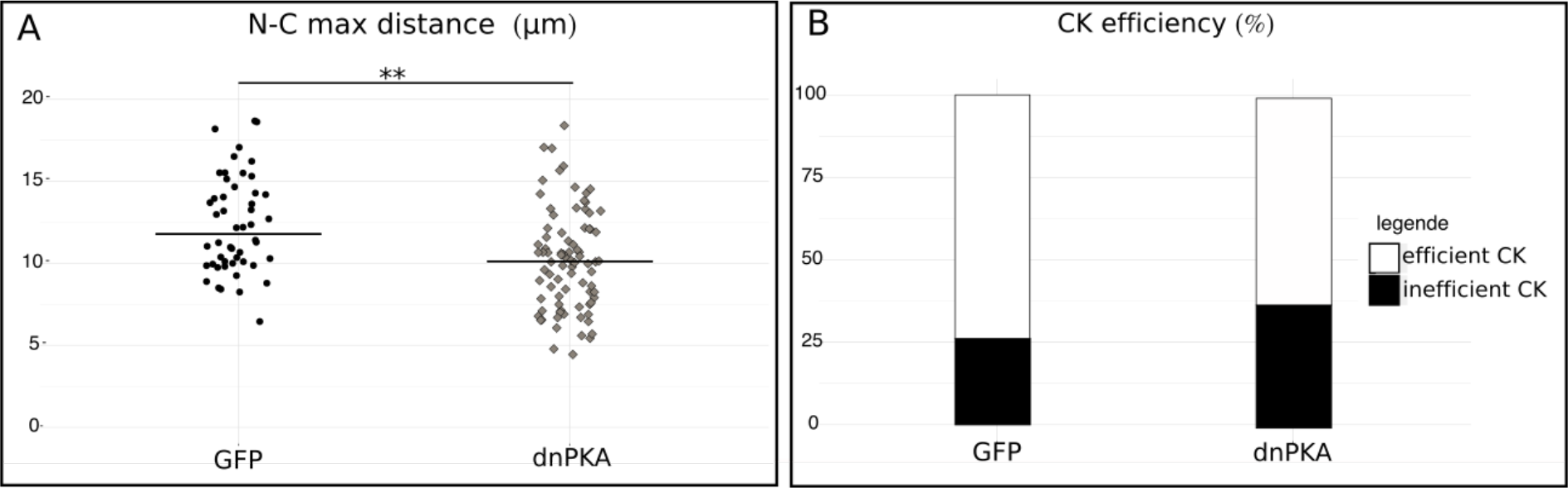
Centrokinesis after PKA delocalization. A) Maximal distance between nucleus and centrosome during CK of neurons electroporated with centrin-RFP and GFP or dnPKA-GFP in Kif3a^lox/lox^ background: the maximal distance was decreased in dnPKA neurons (GFP 12.32±0.42μm versus dnPKA 10.46±0.38μm, One-way Kruskal-Wallis test (Chi-square X^2^=13.54, p=0.003, df=3, followed by Nemenyi test (** p<0.01). The black line represents the median B) Percentage of CK efficiency in neurons electroporated with centrin-RFP and GFP or dnPKA-GFP in Kif3a^lox/lox^ background: the percentage of inefficient CK was increased in dnPKA neurons (GFP 26% versus dnPKA 37%, Pearson’s Chi-squared test X^2^= 6.06, p=0.027) GFP: N=3, n=50, dnPKA: N=3, n=86

**Figure S7:**
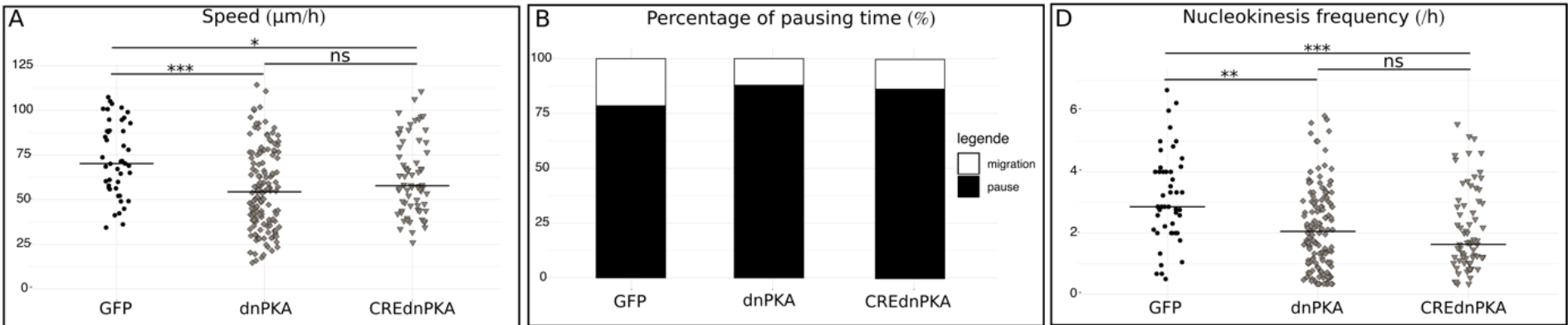
Migration defects after PKA delocalization alone or PKA delocalization along with cilium ablation. The CREdnPKA neurons are not significantly different from the dnPKA neurons for the three parameteres analyzed A) Speed of neurons electroporated with GFP, dnPKA-GFP or dnPKA-GFP plus CRE-TdTomato in Kif3a^lox/lox^ background: GFP 75.49±3.48μm/h versus dnPKA 55.76±1.90μm/h and CREdnPKA 63.73±2.85 μm/h (one-way anova (F_(4,383)_=7.87, p<0.001, followed by Tukey HSD test (* p<0.05, *** p<0.001) B) Percentage of pausing time of neurons electroporated with GFP, dnPKA-GFP or dnPKA-GFP/CRE-TdTomato in Kif3a^lox/lox^ background: 76% in GFP versus dnPKA 86% and CREdnPKA 84% (Pearson’s Chi-squared test X^2^= 67.25, p<0.001) C) Nuclear translocation frequency of neurons electroporated with GFP, dnPKA-GFP or dnPKA-GFP/CRE-TdTomato in Kif3a^lox/lox^ background: GFP 3.15±0.21 NK/h versus dnPKA 2.23±0.12NK/h and CREdnPKA 2.09±0.16NK/h (One-way Kruskal-Wallis test (Chi square X^2^= 19.57, p<0.001, df=4, followed by Nemenyi test (* p<0.05, *** p<0.001) The black line represents the median. GFP: N=3, n=48, dnPKA: N=3, n=146, CREdnPKA: N=3, n=86

**Figure S8:**
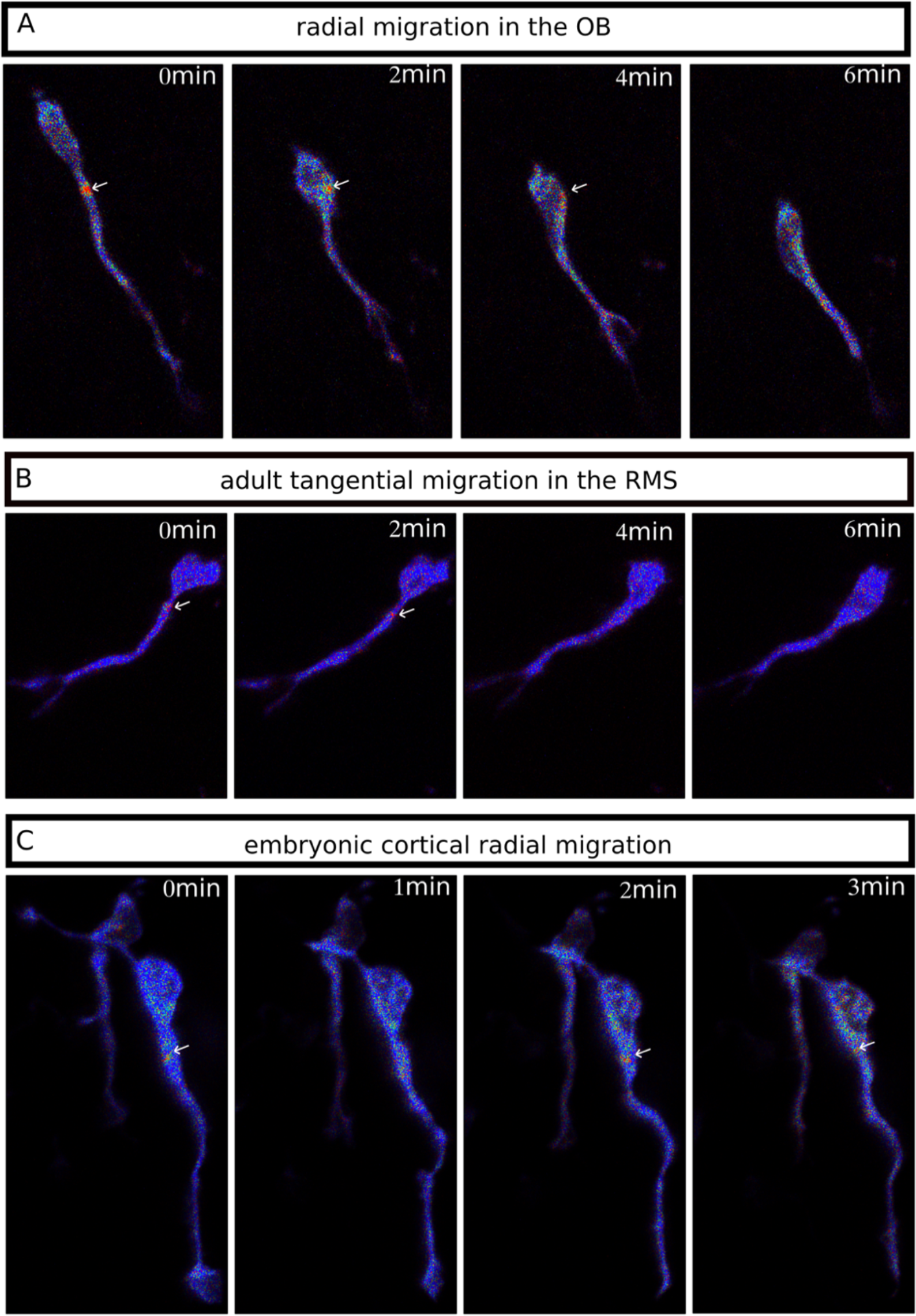
The cAMP hotspot is present in different kind of migrating neurons. A) Live two-photon imaging of a radially migrating neuron in the olfactory bulb transfected by Epac-S^H18^7. The cAMP hotspot is present during NK. B) Live two-photon imaging of a tangentially migrating neuron in the adult RMS transfected by Epac-S^H18^7. The cAMP hotspot is present during NK. C) Live two-photon imaging of a radially migrating neuron in the embryonic cortex transfected by Epac-S^H18^7. The cAMP hotspot is present during NK.

## MATERIAL AND METHODS

### Mouse lines

Mice were housed in a 12-hour light/dark cycle, in cages containing 2 females and one male. The postnatal mice were housed in the cages with their parents. Animal care was conducted in accordance with standard ethical guidelines (National Institutes of Health publication no. 85-23, revised 1985 and European Committee Guidelines on the Care and Use of Laboratory Animals 86/609/EEC). The experiments were approved by the local ethics committee (Comité d’Ethique en Expérimentation Animale Charles Darwin C2EA-05 and the French Ministère de l’Education Nationale de l’Enseignement Supérieur et de la Recherche APAFIS#13624-2018021915046521_v5). We strictly performed this approved procedure. The mice used were in a C57BL6-J background. Kif3a conditional knockout (Kif3a^lox/lox^) (*28*) and Rpgrip1l conditional knockout (Rpgrip1l^lox/lox^) were genotyped according to the original protocols. Kif3a is part of a kinesin motor required for cilium maintenance (*26*). Rpgrip1l is a protein located at the transition zone of the PC, which mutation leads to ciliopathies in humans (*29*).

### Plasmid constructions

pCaggs-Epac-S^H187^ was derived from pCDNA3-Epac-S^H187^ kindly given by Kees Jalink (*14*). Minor modification on the Epac-S^H187^ sequence was performed to reduce codon redundancy without altering the protein sequence. pCaggs-GFP, pCaggs-CRE-IRES-GFP and pCaggs-CRE-IRES-Tdtomato were designed in the laboratory. pCIG-PKA RIΔ2-6 wt subunit flagged (call later on pdn-PKA) was kindly given by Elisa Marti (Saade et al, 2017). pCs2-Centrin-RFP was ordered in addgene (#26753). All the plasmids were used at a concentration between 5 to 8μg/μL (0,01% Fast green) for postnatal electroporation, 2μg/μL (0,01% Fast green) for in utero electroporation and 10μg/μL for adult electroporation.

### Biosensor

The Epac-SH187 cAMP biosensor is composed of a part of Epac protein coupled to a donor and an acceptor fluorophore (13). This biosensor displays a high ratio change and excellent photostability for measuring live cAMP concentration in the micromolar range. It switches from a high FRET conformation to a lower FRET conformation upon binding of cAMP on EPAC. Changes in cAMP concentration were analyzed by ratioing donor (mTurquoise2) and acceptor (td-cpVenus) images, and represented by pseudo-colors (see Fig.1C,D).

### miRNA production

Silencing of AC3 has been performed using BLOCK-iT™ Pol II miR RNAi Expression Vector kits (Invitrogen) and the RNAi Designer (Invitrogen). The sequence of the single stranded oligos are AC3.3: top: TGCTGTTAGGATGGAGCACACGGCATGTTTTG GCCACTGACTGACATGCCGTGCTCCATCCTAA, bottom: CCTGTTAGGATGGAGCACGGCATGTCAGTCAGTGGCCAAAACATGCCGTGTGCTCCATCCTAAC. The double stranded oligos were inserted in a pcDNA™6.2-GW/EmGFP-miR. To produce the pcDNA™6.2-GW/Tdtomato-miR, GFP was replaced by tdTomato using Dra1 and BamHI. The resulting constructions were sequenced before use.

### Postnatal electroporation

Postnatal electroporation was performed at P2. The postnatal mice were anesthetized by hypothermia. Pseudo-stereotaxic injection (from lambda ML: −1,2, A/P:2, D/V: 2,5-2) using glass micropipette (Drummond scientific company, wiretol I 50μL, 5-000-1050) was performed and 2μL of plasmid (betweeen 5 and 8 μg/μL) were injected. Animals were subjected to 5 pulses of 99,99V during 50ms separated by 950ms using the CUY21 SC Electroporator and 10mm tweezer electrode (CUY650-10 Nepagene). The animals were placed on 37°C plates to restore their body temperature before returning with their mother. Animals were considered as fully restored when pups were moving naturally, and their skin color returned to pink.

### Acute brain slices

Brain slices from mice aged from P6 to P10 were prepared as previously described (*30*). Pups were killed by decapitation and the brain was quickly removed from the skull. 250μm sagittal brain slices were cut with a VT1200S microtome (Leica). Slices were prepared in the ice-cold cutting solution of the following composition: 125mM NaCl, 0.4mM CaCl_2_, 1mM MgCl_2_, 1.25mM NaH_2_PO_4_, 26mM NaHCO_3_, 5mM sodium pyruvate, 20mM glucose and 1mM kynurenic acid, saturated with 5% CO_2_ and 95% O_2_. Slices were incubated in this solution for 30 min at room temperature and then placed in recording solution (identical to the solution used for cutting, except that the Ca^2+^ concentration was 2mM and kynurenic acid was absent) for at least 30 min at 32°C before image acquisition.

### Time-lapse video microscopy

To analyze cell migration and centrosome dynamics, images were obtained with an inverted SP5D confocal microscope (Leica). Images were taken every 3 min for 2-3h using a 40X/ 1,25 N.A. objective with 1.5 optical zoom.

Biosensor images were acquired with an upright two-photon microscope Leica SP5 MPII with a 25×/ 0,95 N.A, objective, 4× optical zoom, and GaAsP hybrid detector. The excitation wavelength was set at 850 nm to excite the the mTurquoise2 donor. The two emission wavelengths were acquired simultaneously with filters of 479+/−40 nm and 540+/−50 nm. Image stacks with 2μm intervals were taken every minute for 1h. The presence of tdTomato, indicative of CRE recombinase or miRNA, was assessed with a confocal head. For simultaneous detection of the centrin-RFP and biosensor, two-photon excitation for the biosensor and confocal head for centrin-RFP were alternated. Stacks were spaced by 1μm and acquired every 2 minutes.

The temperature in the microscope chamber was maintained at 32°C, for embryos and postnatal imaging, or 35°C, for P30 imaging, and brain slices were continuously perfused with heated recording solution (see above) saturated with 5% CO2 and 95% O2.

### Analyses of neuronal migration

Analyses were performed using ImageJ (NIH Image; National Institutes of Health, Bethesda, MD) software and MTrackJ plugging (*31*). The nucleus and the centrosome of each cell were tracked manually on each time frame during the whole movie. For cell migration and centrosome movement characteristics, calculation of speed, nuclear translocation frequency and pausing time were performed using the x,y,t coordinates of the nucleus of each cell.

Cells were excluded from the analysis if they were tracked during less than 30 min or did not perform any nuclear translocation during the whole tracking. A cell was considered as migrating if it performed a distance superior to 6 μm during a 3-minute interval.

A centrokinesis was defined as a forward movement superior to 2 μm followed by a backward movement superior to 2 μm. Maximal distance between centrosome and nucleus of every centrokinesis was considered.

### Quantification of biosensor images

Image stacks obtained for donor and acceptor emission were processed with a custom code developed in the IGOR Pro environment (Wavemetrics, Lake Oswego, OR, USA). The maximum intensity was projected vertically to form a 2D image. The fluorescence intensity of donor and acceptor were averaged to build an image indicating biosensor concentration. The fluorescence intensity of donor and acceptor were ratioed for each pixel to report biosensor activation level (see above). For each experiment, ratio values were multiplied by a constant such that baseline ratio was 1. These images were combined to produce pseudocolor images, with the ratio used as hue (from blue to red) and the intensity as value, while the saturation was set to the maximum (30, 31).

For each migrating neuron, the ratio was analyzed in 1 dimension, along its migration direction, and calculated as follows. A series of anchor points were manually positioned along the length of the cell, and the profile of ratio and intensity was calculated along these line segments, over a total width of 1 μm. The intensity profile was annotated manually, marking the rear of the cell and the tip of the leading process of the migrating neuron, and both sides of the nucleus. The movement of the nucleus during the recording was then used to determine the orientation of the migration, by fitting a line to the positions of the nucleus for all time points. Coordinates in the frame of reference of the image were then converted to a single position along the axis of cell migration. The width of the hotspot was the width of a Gaussian function fitted to the ratio trace.

### Immunohistochemistry

P7-P10 mice were lethally anesthetized using euthasol. Intracardiac perfusion with 4% paraformaldehyde were performed. Brain were postfixed overnight in 4% paraformaldehyde. Three rinses were done with PBS 1x (gibco 1400-067). 50μm sagittal slices were cut with VT1200S microtome (Leica). Slices were placed one hour in a saturation solution (10% fetal bovine serum; 0,5% Triton-X in PBS). The primary antibodies used in this study were: GFP (Aves, GFP-1020, 1/1000), PKAc (Cell Signaling Technology, #4782 1/250), AC3 (Santa Cruz, C-20 sc-588, 1/200), Arl13b (UC Davis/NIH NeuroMab Facility, 75-287, 1/1000), ɣ-tubulin (Sigma-Aldrich, T6557, 1/500). The antibodies were diluted in saturation solution. Slices were incubated 48 to 72h at 4°C under agitation with the antibodies. Three rinses were done with PBS 1x. The secondary antibodies used were: anti-chicken IgY alexa Fluor 488 (1/1000, Jackson ImmunoResearch: 703-545-155) against anti-GFP, anti-rabbit IgG Cy5 (1/1000, Jackson ImmunoResearch: 711-175-152) against anti-PKAc, anti-rabbit IgG Cy3 (1/2000, Jackson ImmunoResearch: 711-165-152) against anti-AC3, anti-Mouse IgG, Fcɣ Subclass 1 specific alexa Fluor 594 (1/2000, Jackson ImmunoResearch: 115-585-205) against anti-ɣ-tubulin, anti-Mouse IgG, Fcγ subclass-2a specific alexa Fluor 647 (1/1000, Jackson ImmunoResearch: 115-605-206) against anti-Arl13b. The antibodies were diluted in saturation solution. Slices were incubated 1h at room temperature under agitation with the secondary antibody solution. Three rinses with PBS 1X were done. Slices were counter-colored with Hoeschst and mounted with Mowiol.

### Statistics

All manipulations and statistical analyses were implemented with R (3.5.1). Normality in the variable distributions was assessed by the Shapiro-Wilk test. Furthermore, the Levene test was performed to probe homogeneity of variances across groups. Variables that failed the Shapiro-Wilk or the Levene test were analyzed with nonparametric statistics using the one-way Kruskal–Wallis analysis of variance on ranks followed by Nemenyi test post hoc and Mann–Whitney rank sum tests for pair-wise multiple comparisons. Variables that passed the normality test were analyzed by means of one-way ANOVA followed by Tukey post hoc test for multiple comparisons or by Student’s t test for comparing two groups. Categorical variables were compared using Pearson’s Chi-squared test or Fisher’s exact test. All the statistical analyses have been performed on the five groups together (GFP, Cre, miRAC3, dnPKA, dnPKA/CRE).

A p value of < 0.05 was used as a cutoff for statistical significance. Results are presented as the mean ± SD or medians and the given values in the text are the mean ± SEM, unless otherwise stated. The statistical tests are described in each Fig.ure legend.

### In utero electroporation

Timed pregnant E15 C57Bl6 mice were anesthetized with isoflurane. During the experiment, mice were on a 37°C plate. The uterine horns were exposed, and the embryos were injected in the lateral ventricle with a glass micropipette. 1μL of plasmid (2μg/μL) was injected. The successfully injected animals were then subjected to 5pulses of 45V during 50ms separated by 950ms using the CUY21 SC Electroporator and 5mm tweezer electrode (CUY650-5 Nepagene). The uterine horns were then replaced in the belly and the belly was sewn up. The animals were placed in a cage on 37°C plates to restore their body temperature before returning in their own cage.

### Adult electroporation

P21 C57Bl6 animals were anesthetized with a mixture of ketamine and xylazine. Stereotaxic injection (from the bregma M/L: 1, A/P:0, D/V:2,1) was performed with a glass micropipette and 2μL of the plasmids (10μg/μL) was injected in the right lateral ventricle. The animals were then removed from the stereotaxic setting and subject to 5 pulses of 200V during 50ms separated by 950ms using the electroporator NEPA21 with 10mm tweezer electrode (CUY650-10). The animals were then placed on a 37°C plates to restore their body temperature before returning in their cage.

### Acute brain slices of embryonic and adult brains

For imaging of E18 embryos, the mother was killed by cervical dislocation and the embryos were removed from the horn and placed in an ice-cold cutting solution (same composition as above). The embryos were decapitated, and the brain was quickly removed from the skull. 300μm sagittal brain slices were cut in the same ice-cold solution. Then the same protocol as above was followed. P30 mice were killed by cervical dislocation. The brain was quickly removed from the skull. 200μm sagittal brain slices were cut with a VT1200S microtome (Leica). Slices were prepared in an ice-cold solution of the following composition: 130mM potassium gluconate, 15mM KCl, 2mM EGTA, 20mM HEPES, 25mM glucose, 1mM CaCl_2_ and 6mM MgCl_2_, supplemented with 0.05mM D-APV (304 mOsm, pH 7.4 after equilibration) saturated with 5% CO_2_ and 95% O_2_ (*32*). Then the slices were places for 3 to 5 minutes in a ice-cold solution of the following composition: 225mM D-mannitol, 2.5mM KCl, 1.25mM NaH2PO4, 25mM NaHCO3, 25mM glucose, 1mM CaCl_2_ and 6mM MgCl_2_ (*32*) saturated with 5% CO_2_ and 95% O_2_ before being placed in the recording solution (described above) at 32°C and saturated with 5% CO_2_ and 95% O_2_ for at least 30 min.

## Acknowledgments

We thank Sylvie Schneider-Maunoury for Rpgrip1l mice. We are grateful to Kees Jalink for Epac SH187 plasmid and to Elisa Marti for dnPKA plasmid. We thank all re-readers including Isabelle Dusart, Stéphane Nédelec, Fiona Doetsch, Nathalie Spassky and Jonathan Weitzman. The experiments were performed in IBPS imaging facility and the mice were housed in IBPS animal facility. We thank Claire Fournier-Thibault for lending her electroporator.

## Funding

this work was supported by CNRS, INSERM, Fondation Lejeune and Sorbonne University. JS was funded by a doctoral fellowship awarded by IBPS to AT and PV labs. This work was supported (in part) by the Investissements d’Avenir program managed by the ANR under reference ANR-11-IDEX-0004-02. AT and PV labs are members of the Bio-Psy Labex.

## Author contributions

JS performed experiments, developed the methodology, analyzed data and wrote the manuscript; MC performed experiments and analyzed data; CD and CF performed experiments; MD performed formal analysis; CM participated in the initiation of the work and provided materials; AT acquired fundings and edited the manuscript; PV acquired fundings, developed software, supervized project and edited the manuscript; IC acquired fundings, performed experiments, conceptualized and supervized the project and wrote the manuscript.

## Competing interests

authors declare no competing interests

## Data and materials availability

all data is available in the main text or the supplementary materials.

## Notes

#### Summary of Updates

A more concise version

